# Quantifying phase separation property of chromatin associated proteins under physiological conditions using anti-1,6-hexanediol index

**DOI:** 10.1101/2020.12.07.415489

**Authors:** Minglei Shi, Kaiqiang You, Taoyu Chen, Chao Hou, Zhengyu Liang, Mingwei Liu, Jifeng Wang, Taotao Wei, Jun Qin, Yang Chen, Michael Q. Zhang, Tingting Li

## Abstract

**Background:** Liquid–liquid phase separation (LLPS) is an important organizing principle for biomolecular condensation and chromosome compartmentalization. However, while many proteins have been reported to undergo LLPS, quantitative and global analysis of chromatin LLPS property remains absent.

**Results:** Here, by combing chromatin associated protein pull-down, quantitative proteomics and 1,6-hexanediol treatment, we developed Hi-MS and defined anti-1,6-HD index of chromatin-associated proteins (AICAP) to quantitative measurement of LLPS property of chromatin-associated proteins in their endogenous state and physiological abundance. The AICAP values were verified by previously reported experiments and were reproducible across different MS platforms. Moreover, the AICAP values were highly correlate with protein functions. Proteins act in active/regulatory biological process often exhibit low AICAP values, while proteins act in structural and repressed biological process often exhibit high AICAP values. We further revealed that chromatin organization changes more in compartment A than B, and the changes in chromatin organization at various levels, including compartments, TADs and loops are highly correlated to the LLPS properties of their neighbor nuclear condensates.

**Conclusions:** Our work provided the first global quantitative measurement of LLPS properties of chromatin-associated proteins and higher-order chromatin structure, and demonstrate that the active/regulatory chromatin components, both protein (trans) and DNA (cis), exhibit more hydrophobicity-dependent LLPS properties than the repressed/structural chromatin components.

## Background

The nucleus of eukaryotic cells is filled with three-dimensionally folded DNA as well as a considerable number of proteins and RNAs, which together form nuclear condensates that regulate gene expression. By using Hi-C technique to analyze chromatin organization, chromatin is partitioned into two compartments termed ‘A’ and ‘B’, which correspond to euchromatin (A) and heterochromatin (B)[1]. While the presence of chromatin compartments is now well established, the mechanisms that drive chromatin condensation and compartmentalization remain largely unclear[2]. Recently, LLPS has been proposed as one of possible mechanisms to explain chromosome compartmentalization[3, 4]. Both euchromatin- and heterochromatin- associated proteins have been shown to be phase-separated, including RNA polymerase II[5, 6], mediator complex subunits[7], heterochromatin proteins 1 (HP1)[8, 9], and Polycomb protein chromobox 2 (CBX2)[10, 11].

However, identifying LLPSs in cells remains challenging due to the limited arsenal of tools. LLPS proteins are commonly identified by low-throughput methods such as droplet roundness/fusion, immunofluorescence and fluorescence redistribution after photobleaching (FRAP)[12]. As a consequence, it is difficult to compare the LLPS properties of different proteins in these compartments, although molecular compositions and biological functions of these spatially segregated chromatin compartments differ considerably. In addition, while LLPS is now believed to be essential for chromosome compartmentalization, DNA loop-extrusion is believed to mediate TAD/loop formation and antagonize to LLPS[13]. Since TAD and loop are subcomponents of chromatin compartment, LLPS may also affect the TAD and loop structure. However, it remains unclear how LLPS functions in specific TAD/loop and cis-regulatory DNA regions. Therefore, it is critical to quantitatively and globally measure the LLPS properties of nuclear condensates and chromatin organization especially in their endogenous abundance.

The aliphatic alcohol 1,6-HD is widely used for disrupting LLPS condensates. It contains a hydrophobic group that is composed of 6 hydrogenated carbon atoms, which interfere with the hydrophobic interactions, and consequently affect hydrophobicity-dependent LLPS condensates[14, 15]. Furthermore, compared with another commonly used detergent, SDS, which consists of twelve hydrogenated carbon atoms, 1,6-HD is sufficiently weak to only disrupt LLPS condensates, without affecting either solid-like assemblies[16] or membrane-bounded organelles[17]. Moreover, a recent study reported that following 1,6-HD treatment of MCF-7 cells, the ChIP-seq peak signal for the transcription factor (TF) GATA3 was significantly weakened, while the peak signal for another TF ER remained unchanged[18], suggesting that chromatin binding proteins exhibit protein-specific sensitivities to 1,6-HD treatment, thus affecting their ability to bind to DNA. This protein-specific sensitivity provides a valuable opportunity to quantitatively measure LLPS properties of nuclear condensates in their endogenous state and physiological abundance.

However, capturing of LLPS-mediated chromatin-associated proteins is difficult. LLPS proteins such as transcription factors or mediators are expressed at lower levels compared with constitutive proteins such as histones. Furthermore, LLPS proteins may bind to DNA with low affinity due to the highly dynamic nature of this interaction[19]. In addition, the methods available for genome-wide capturing chromatin-associated proteins are too hash to capture LLPS proteins. For instance, for CHEP, cells are usually washed using 4% SDS and 8M urea[20]; while for DEMAC, high speed centrifugation for at least 48h is needed[21]. Thus, new methods that capable of effectively capture LLPS-mediated chromatin-associated proteins are urgently required.

Here, we developed a method called Hi-MS to quantitatively and globally measures the LLPS properties of chromatin-associated proteins by combining chromatin-associated protein pull-down, quantitative proteomics and 1,6-HD treatment. We also analyzed how LLPS affects chromatin organization combined with Hi-C after 1,6-HD treatment. By applying these methods, we obtained a first global view of LLPS properties of nuclear condensates and chromatin organization in their endogenous state.

## Results

### Development and validation of Hi-MS

In order to quantify chromatin-associated protein changes after 1,6-HD treatment, we developed a method that effectively captures chromatin-associated proteins in situ (Figure 1A). To enrich regulatory proteins, we targeted gene promoter regions based on genome sequence preference. As shown in Figure 1B, GGCC is a nucleotide sequence enriched in gene promoter regions. We previously developed a method called BL-Hi-C[22], which uses restriction endonuclease *Hae*III to cut at GGCC sites to enrich cis-regulatory elements of gene promoters, including both activated elements marked by H3K27Ac and repressed elements marked by EZH2 (Figure 1C). This procedure is relatively gentle, and most of the chromatin-associated proteins are well preserved. Here, we used a protocol based on BL-Hi-C to extract chromatin-associated proteins. Briefly, we crosslinked cells using 1% formaldehyde and then digested the genome by *Hae*III, then the digested DNA fragment ends were ligated via biotinylated bridge linker. Next, we sonicated the cells, and the biotinylated linker/DNA/protein complexes were captured by magnetic streptavidin-beads, before label-free quantitative mass spectrometry (MS) analysis. We named this method Hi-MS corresponding to the name of Hi-C (Figure 1A). We used Hi-MS to extract chromatin-associated proteins in K562 cell line, which is a widely used cell line with an extensive public omics data set. Typically, 10^8^ cells are required for genome-wide chromatin protein capture methods[20, 21], while for Hi-MS analysis, 10^7^ cells are sufficient. Importantly, TFs and cofactors were enriched 5-fold in the Hi-MS sample compared to undigested control samples. Other nuclear proteins involved in mRNA processing, transcription, DNA repair and chromosome organization were enriched 4-15 folds, while proteins involved in cytoskeleton organization represented by KRTs were significantly reduced (Figure 1D).

**Figure 1.**
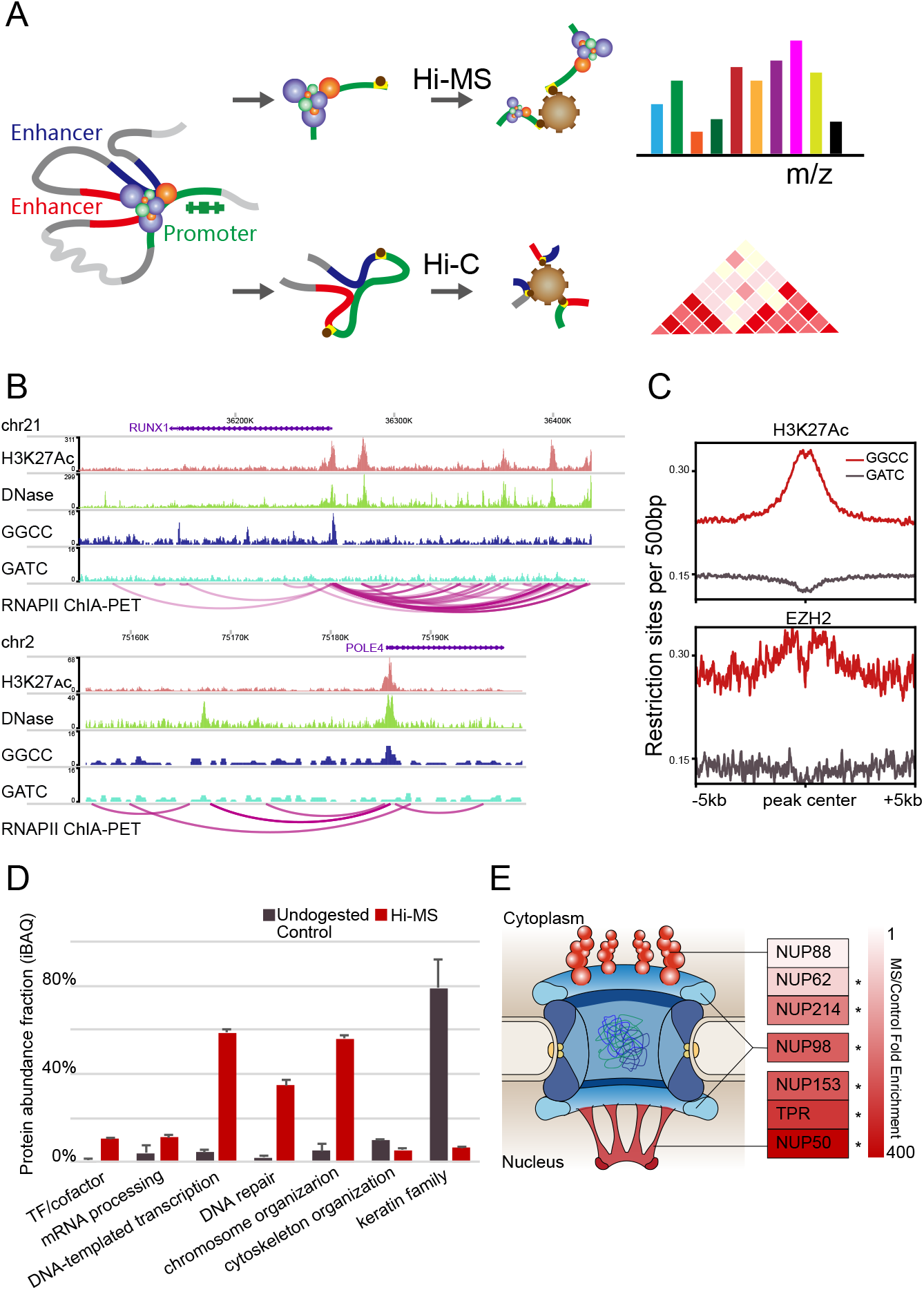
Hi-MS effectively enrich chromatin-associated proteins. **A.** Schematic of Hi-MS. **B.** The examples of GGCC distribution in the gene promoter region. Top: RUNX1; bottom: POLE4. These two regions were selected as represents of super enhancer and typical enhancer, according to (Hnisz et al., 2017). **C.** The distribution of human genome sequence GATC and GGCC proximate to H3K27Ac and EZH2 binding peaks. **D.** Chromatin associated proteins are effectively enriched by Hi-MS (MS) compared with mock (ML). **E.** MS/Control fold enrichment of NUPs agreed with their spatial location. NPC model was adapted from (Buchwalter et al., 2019).

Next, we determined the efficiency of our Hi-MS method in enriching chromatin-associated proteins by analyzing the components of nuclear pore complex (NPC). As shown in Figure 1E, cytoplasmic filament components, which locate on the cytoplasm side of NPC, including NUP214, NUP88 and NUP62, showed an average fold enrichment of 102. In comparison, nuclear basket components, which locate on the nuclear side of NPC, including TPR, NUP50 and NUP153, showed an average fold enrichment of 339. The nuclear/cytoplasmic ring component NUP98, which locates in the middle of the NPC, showed a fold enrichment of 262. Together, these results indicated that Hi-MS is a sensitive and efficient method to capture chromosome-associated proteins *in situ*.

### Proteins exhibit different sensitivities to 1,6-HD treatment

In order to effectively measure the sensitivity of chromatin-associated proteins to 1,6-HD treatment by quantitative proteomics, we first titrated the concentration of 1,6-HD by testing local distribution of several proteins using immunofluorescence. It was previously reported that in HeLa cells, stress granules (SG) and P bodies (PB) can be dissolved following 1,6-HD treatment[23]. We based our incubation time on this method, and tested the dissolution of MED1/FUS puncta at different concentrations of 1,6-HD in K562 cells. As shown in Additional file 1: Figure S1A-B, 10% 1,6-HD effectively dissolved most MED1 puncta. We failed to detect any FUS puncta; however, with increasing 1,6-HD concentrations, the evenly distributed FUS gradually translocated into the cytoplasm. As 1,6-HD has the capability to dissolve the channel in NPC[24], we speculate that this translocation occurred because the NPC was destroyed.

Kroschwald et al. also found that after the dissolution of stress granules, smaller stress granule-like structures re-accumulate in a significant percentage of cells. However, hypotonic medium effectively alleviates this re-accumulation[16], probably because cells swell slightly in hypotonic conditions, thus reducing the local protein concentration. Based on this report, we mixed 30% 1,6-HD aqueous solution with normal K562 cell culture medium, and obtained a 2/3 dilution of medium with 10% 1,6-HD. To evaluate the sensitivity of proteins to this hypotonic 1,6-HD treatment, we used the cytoplasmic/nuclear proteins ratio as a measurement of sensitivity to 1,6-HD treatment. As shown in Additional file 1: Figure S1C-D, the order of sensitivity was MED1> FUS > EZH2 > H3.

Together, these results indicated that each protein is characterized by a protein-specific sensitivity to 1,6-HD treatment for their interaction with genomic DNA/retention within the nucleus, thus allowing for subsequent quantitative measurement of this sensitivity using Hi-MS.

### Evaluating LLPS properties of proteins using AICAP

In this section, we set out to quantitatively measure the sensitivity of chromatin-associated proteins to 1,6-HD treatment by Hi-MS. As demonstrated earlier, proteins exhibit different sensitivities to 1,6-HD treatment. With the quantified protein amount before and after 1,6-HD treatment, we defined an anti-1,6-HD index of chromatin-associated proteins (AICAP) (Figure 2A). This index quantitatively reflects the sensitivity of every chromatin-associated proteins to 1,6-HD treatment.

**Figure 2.**
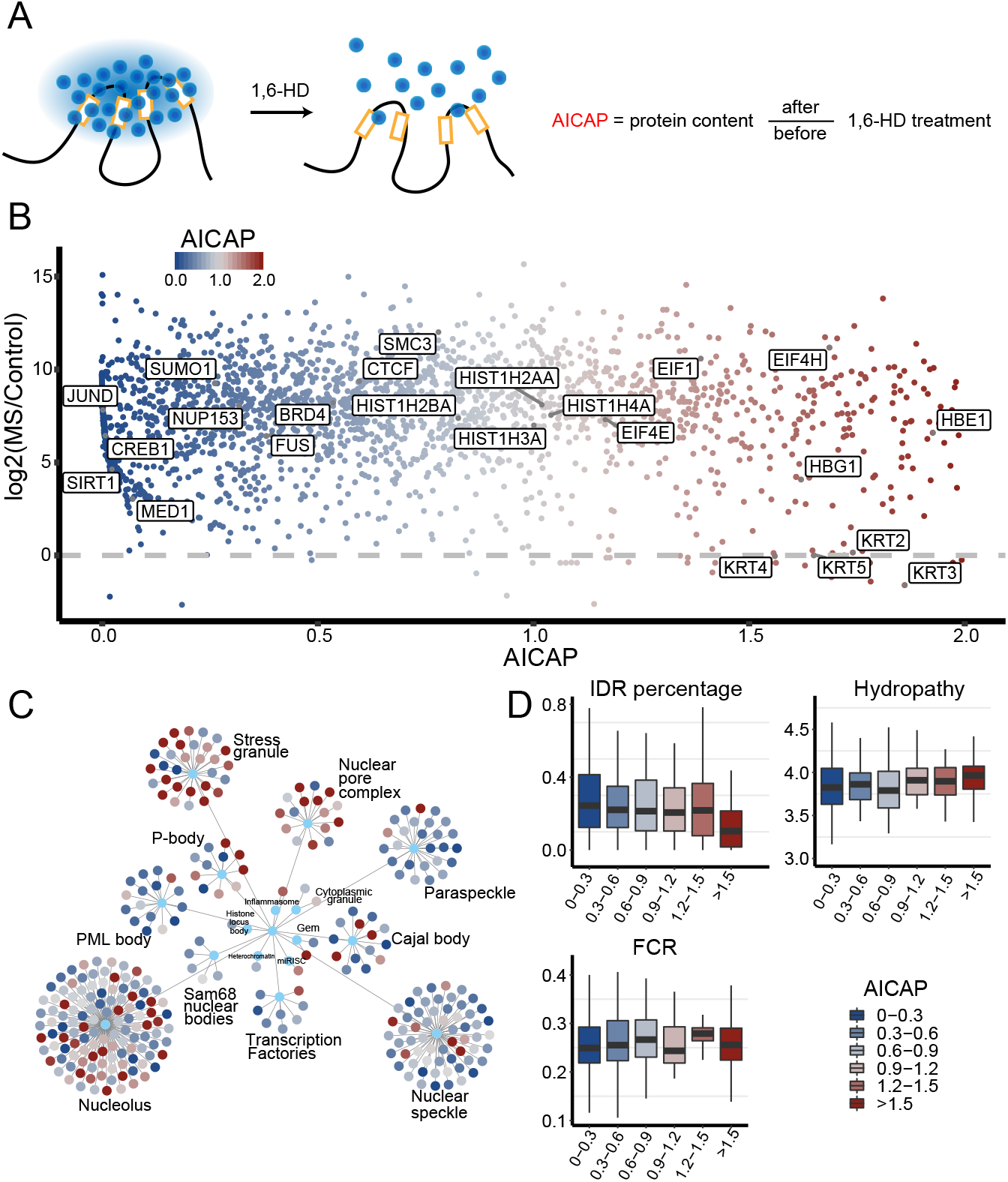
AICAP represented hydrophobicity-dependent LLPS properties. **A.** Schematic of 1,6-HD treatment and definition of AICAP. **B.** Scatter plot of proteins with AICAP values 0~2 captured by Hi-MS. **C.** AICAP values of proteins in nuclear domains. Proteins of every nuclear domain were extracted from PhaSepDB (You et al., 2020). **D.** The intrinsic sequence compositions and physical properties of 6 groups of proteins. IDR, intrinsically disorder region; FCR, fraction of charged residue.

We prepared three batches of biological duplication samples using Hi-MS, each batch containing three treatments, 1,6-HD−, 1,6-HD+ and undigested control. As shown in Additional file 1: Figure S2A, the same treatment in different biological duplications can be well clustered, indicating high reproducibility of 1,6-HD treatment. We obtained the AICAP values for 3228 chromatin-associated proteins through mass spectrometry (MS) analysis (Additional file 2: Table S1). Lower AICAP values indicate that the corresponding proteins are more sensitive to 1,6-HD treatment. As shown in Figure 2B, proteins that can undergo LLPS such as FUS, SUMO1, MED1 and YY1 showed low AICAP values; the chromatin architecture protein CTCF, SMC3 showed relatively high AICAP values but still lower than 1.0; proteins of the histone family showed AICAP values around 1.0; while KRTs, EIFs and other typical cytoplasmic proteins showed AICAP values above 1.0. We also displayed AICAP of proteins in the main membrane-free organelles (Figure 2C). The result showed that most proteins in nuclear domains such as nuclear speckles, paraspeckles and PML bodies exhibit low AICAP values, which is in agreement with a previously published LLPS database[25].

Next, we analyzed the intrinsic sequence compositions of captured proteins. Proteins with a high fraction of charged residue (FCR) may interact with each other more via electrostatic forces, while proteins with high hydropathy may do so more via hydrophobic forces. We divided Hi-MS captured proteins into 6 groups based on their AICAP. Our results showed that the AICAP 0-0.3 group exhibited the highest intrinsic disorder region (IDR) percentage and low fraction of charged residue (FCR) content (Figure 2D), which indicates that DNA binding of these proteins depends more on hydrophobicity-dependent LLPS. The AICAP 0.6-0.9 group exhibited the lowest hydropathy but high FCR content. These data suggested that DNA binding of proteins in this group, such as EZH2, RING1, CBX5 depends more on the electrostatic interactions and less on hydrophobicity, and is therefore less sensitive to 1,6-HD treatment. The AICAP 1.2-1.5 group also exhibited high FCR content. This part of the protein may contact DNA non-physiologically. As DNA is negatively charged, it is very likely that proteins containing more positive charges would translocate into the nucleus and bind DNA with more abundance after NPC was destroyed.

In summary, these results indicated that our AICAP provides valuable information for evaluating how DNA binding of chromatin-associated proteins depends on hydrophobic or electrostatic forces.

### AICAP values were verified by previously reported experiments in cell and in vitro

To verify the reliability of AICAP, we compared the AICAP values of proteins with previously published data[7]. They compared the ChIP-seq data of BRD4, MED1 and RNAPII before and after 1,6-HD treatment in mESC cells. The result showed that at the super enhance (SE) region of Klf4, the occupancy levels of BRD4, MED1 and RNAPII were reduced by 44%, 80%, and 56%, respectively. We reprocessed their ChIP-seq data, and calculated the reads density at all SEs before and after 1,6-HD treatment, and found that the occupancy levels of BRD4, MED1 and RNAPII were reduced by 20%, 38.4%, and 21.4%, respectively, which is similar to that of Klf4 (Figure 3A). The results obtained from that ChIP-seq data were consistent with our AICAP results, with the AICAP values of 0.537 for BRD4, 0.072 for MED1 and 0.401 for RNAPII. Sabari et al. also quantify the LLPS properties of MED1 and BRD4 using droplet formation and FRAP assays and found considerable difference between MED1 and BRD4. In FRAP analysis, they found that mEGFP-BRD4 and mEGFP-MED1 puncta recovered with apparent diffusion coefficients of ~0.37 ± 0.13 and ~0.14 ± 0.04 mm^2^/s, respectively. In droplet disturbing assays, the droplet size of MED1-IDR decreased more than that of BRD4-IDR with increasing NaCl concentration. These differences between MED1 and BRD4 agreed with their AICAP values, which further proved the reliability of AICAP[7].

**Figure 3.**
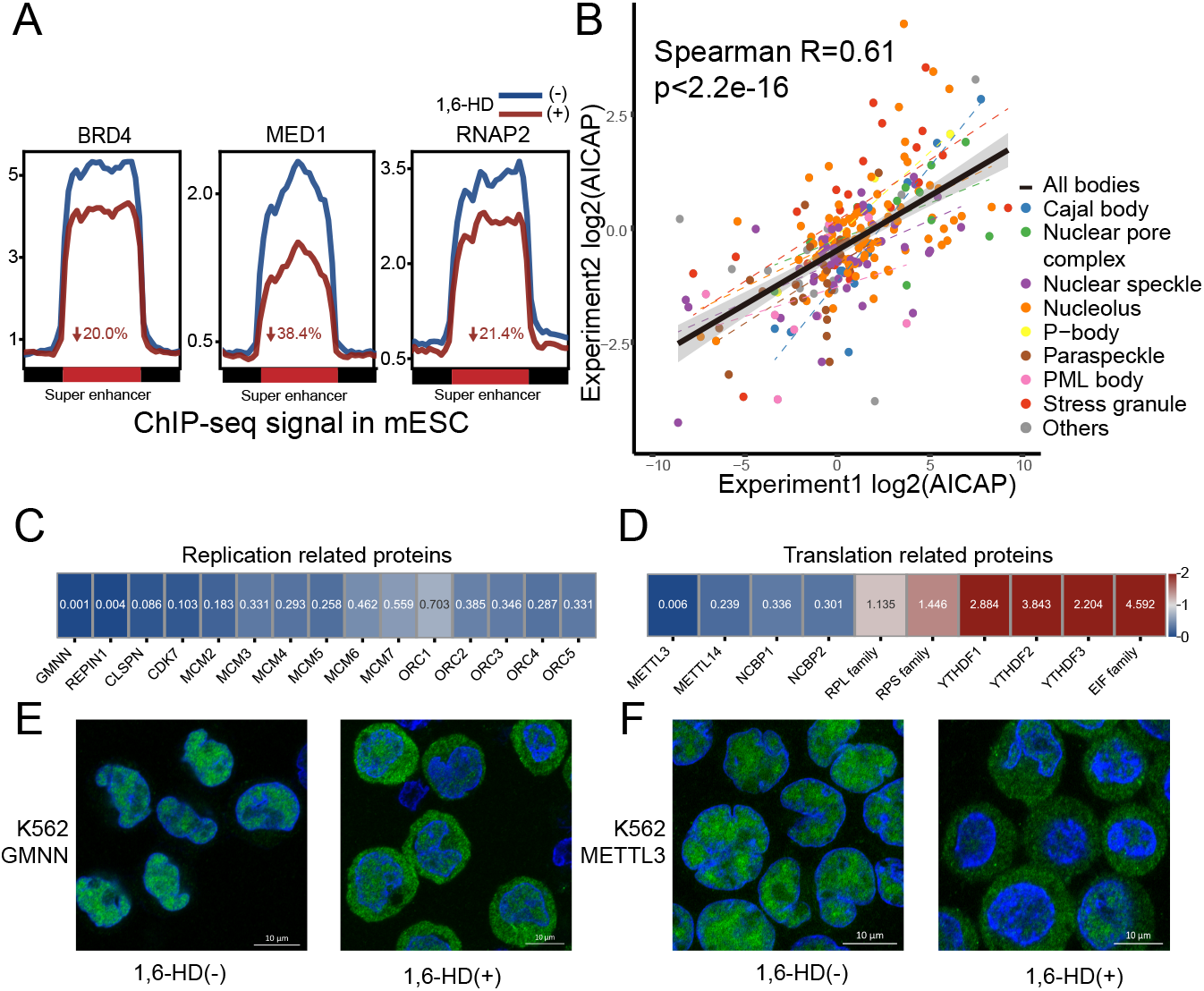
Validity and repeatability of AICAP. **A.** ChIP-seq signal enrichment of BRD4, MED1 and RNAPII at regions defined as super enhancer in mESC before (1,6-HD−) and after (1,6-HD+)1,6-HD treatment. The decreased percentage of signal are noted. Data resource (Sabari et al., 2018). **B.** Repeatability of AICAP values of proteins in nuclear domains between two independent replicates. **C** and **D.** AICAP of DNA replication and translation associated proteins. **E** and **F.** Immunofluorescence of GMNN and METTL3 before (−) and after (+) 1,6-HD treatment.

### AICAP values were reproducible across different MS platforms

To test the robustness of our AICAP, we treated different batches of cultured cells, performed different types of digestion (in gel or in solution) and analyzed on different types of mass spectrometers (Q Exactive or Orbitrap Fusion Lumos Plus). As shown in Additional file 1: Figure S2B, the AICAPs generated by the two batches of experiments significantly correlated with each other. The Spearman correlation coefficient of two batches reached 0.533 with the p-value 1.5e-145. More proteins were obtained using in solution digestion (condition 2 in methods), so the MS data used in following analysis were form condition 2. We further compared the AICAP of common nuclear condensates proteins, and found that the AICAP of proteins in the same condensate correlated well. The Spearman correlation coefficient of nuclear condensates proteins between two batches reached 0.61, with a p-value of 2.2e-16 (Figure 3B). Together, these results demonstrated that the AICAP values generated by our Hi-MS method were robust.

### AICAP values provide a ranked list of LLPS protein candidates

By quantitatively measuring the sensitivity of chromatin-associated proteins to 1,6-HD treatment, we obtained AICAP values for thousands of proteins. In addition to known LLPS proteins, our AICAP values revealed a considerable number of novel LLPS candidates. For example, two components of the DNA replication initiation complex, namely ORC1-5 and MCM 2-7, exhibited low AICAP values (Figure 3C). Moreover, GMNN, CLSPN, REPIN1 and CDC7, all of which play important functions during DNA replication initiation, exhibited low AICAP values (Figure 3C). In translation-related processes, N6-methyladenosine (m6A) RNA modification has been reported to facilitate LLPS of YTHDF [26–28]. Here, we found that the m6A methyltransferase METTL3/14 was highly sensitive to 1,6-HD treatment (Figure 3D). In particular, METTL3, the m6A methyltransferase catalytic subunit, exhibited an AICAP value as low as 0.006, while METTL14, the non-catalytic subunit, exhibited an AICAP value of 0.239. To identify the LLPS properties of GMNN and METTL3, we performed immunofluorescence experiments. As shown in Figure 3E and 3F, both proteins formed clusters in the K562 nucleus. Upon 1,6-HD treatment, these two factors translocated into the cytoplasm, similar to FUS and MED1. Additional file 2: Table S1 provides a complete list of candidate LLPS proteins for further studies.

### Global Reorganization of the 3D Genome after 1,6-HD treatment

Following 1,6-HD treatment, chromatin-associated proteins dissociate from the chromatin at various amounts. To study the effect of this dissociation on the higher-order chromatin organization, we constructed BL-Hi-C libraries in untreated K562 cells or flowing 1,6-HD treatment. For each treatment, we constructed three biological replicates. As shown in Additional file 1: Figure S3A, both libraries clustered well, indicating that the effect of 1,6-HD on chromatin organization is highly reproducible.

Next, we checked the global changes for DNA interactions. As shown in Additional file 1: Figure S3B and S3C, the short-range interactions decreased upon 1,6-HD treatment, while long-range interactions increased. In addition, the trans% interaction reads ratio, which is generally considered to represent the noise ratio, was 17% for the BL-Hi-C library, and 23% after 1,6-HD treatment (Additional file 1: Figure S3D, Additional file 3: Table S2). Compared with a recently published K562 liquid chromatin Hi-C data, the trans% ratio of the Hi-C library was 53%. Following *Dpn*II pre-digestion, this ratio reached 80% (Additional file 1: Figure S3D)[29]. These results further proved that BL-Hi-C is capable of detecting the cis-unique DNA interactions with higher efficiency.

### Chromatin/proteins in active transcriptional regions are more sensitive to 1,6 HD treatment

Using BL-Hi-C data, we tested the DNA-DNA interaction changes of functional DNA elements to explore their sensitivity to 1,6-HD treatment. Functional DNA elements were characterized by 15 types of Epigenome chromatin States (Roadmap Epigenomics Consortium, Nature 2015). As shown in Figure 4A, the intra-chromosome interaction of active transcription regions, including active TSSs and enhancers were most affected, while repressed chromatin regions/areas, including bivalent regions and heterochromatin were less affected. Moreover, the AICAP of hallmark proteins in these regions showed good consistency with the sensitivity to 1,6-HD treatment of these functional DNA elements (Figure 4B). Transcription activation and regulatory proteins exhibit low AICAP values, while heterochromatin proteins and chromatin structural proteins exhibit high AICAP values.

**Figure 4.**
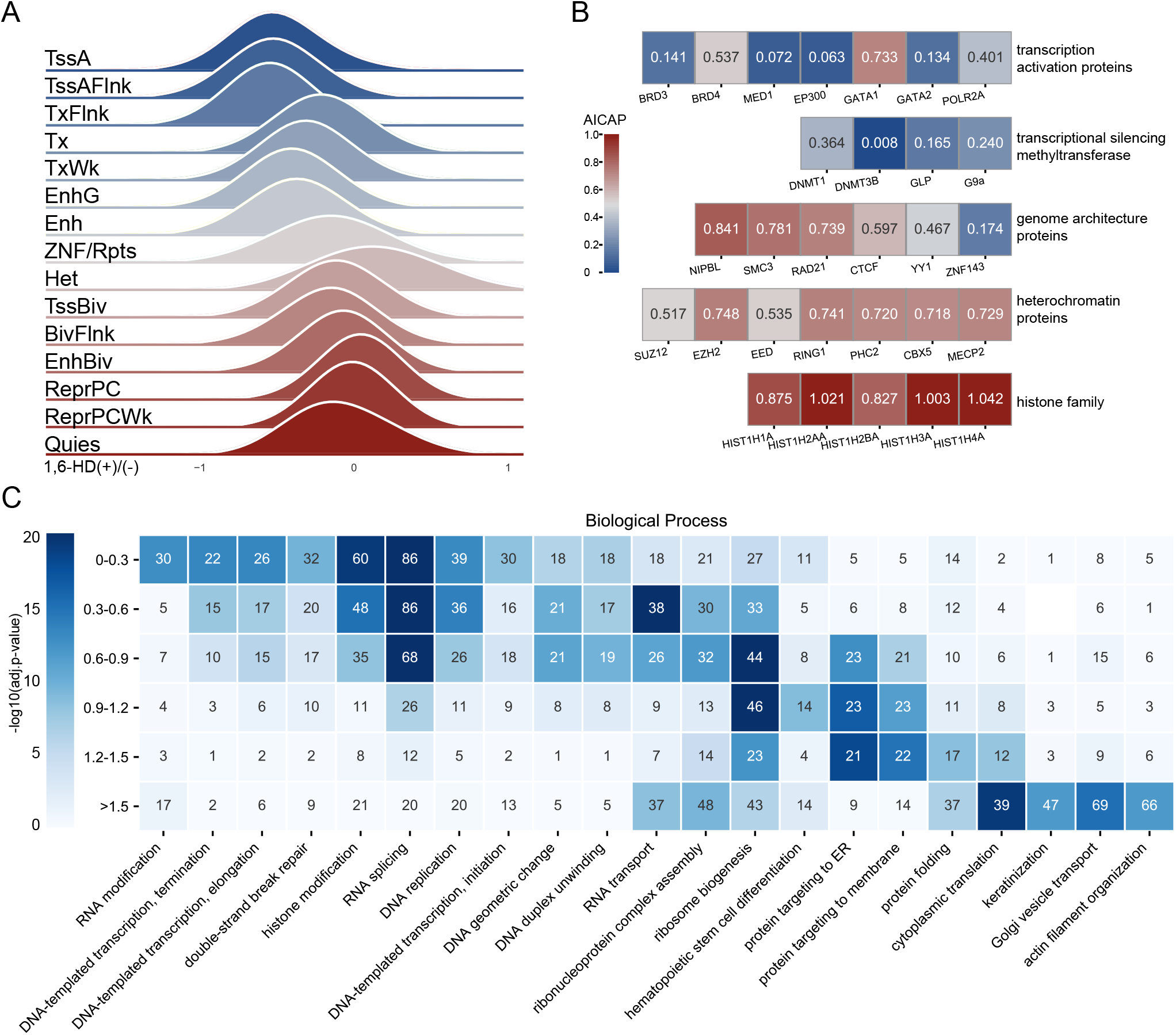
Chromatin/proteins in active transcriptional regions are more sensitive to 1,6 HD treatment. **A.** Intra-chromosome interactions of 15 types epigenome chromatin states after 1,6-HD treatment. **B.** The AICAP values of hallmark proteins in different functional categories. **C.** Gene ontology biological process enrichment analysis of proteins. Proteins were divided into 6 groups based on AICAP(Y-axis). Number of proteins was noted in the corresponding cell.

Apart from these hallmark proteins, the AICAP values of all TFs and cofactors captured by Hi-MS also correlated well with protein function (Additional file 4: Table S3). When we divided all TFs and cofactors into six groups based on AICAP, the ‘biological process’ (BP) terms enriched in group ‘0-0.3’ mainly constituted highly dynamic processes, such as signaling pathway and transcription initiation. The BP terms enriched in group ‘0.3-0.6’ mainly constituted chromatin remodeling, chromatin silencing and other structural and inhibitory processes (Additional file 1: Figure S4A).

Finally, we studied the relationship between AICAP and the functions of all proteins captured by Hi-MS (Additional file 5: Table S4). We divided these proteins into 6 groups based on their AICAP values, and proteins in each group were clustered according to categories of ‘biological process’ (BP, Figure 4C), ‘molecular function’ (MF, Additional file 1: Figure S4B), or ‘cellular component’ (CC, Additional file 1: Figure S4C). As shown in Figure 3C, the main enrichment BP terms within group “0-0.3” constituted RNA modification/splicing, DNA transcription, histone modification and the corresponding CC terms of RNA pol II/TF complex, nuclear speckle, all of which were active regulatory processes. The enriched BP terms near the AICAP value of 1.0 were primarily constitutive nuclear condensate represented by ribosomes, whose main components (RPLs, RPSs) shuttle between the cytoplasm and nucleoli, and frequently contact DNA[30]. Proteins with AICAP values above 1.0 were related to protein translation, folding, and transport, most of which are usually located in the cytoplasm.

Together, the results strongly suggest that our AICAP index highly correlates with protein function. Proteins involved in dynamically regulated processes commonly exhibited low AICAP values. In contrast, proteins involved in stable structural processes often exhibited high AICAP values.

### Chromatin compartment changes are related to neighboring nuclear condensates

Here, we studied the Protein/DNA sensitivities to 1,6-HD treatment in the scope of 3D chromatin structure. 3D chromatin forming hierarchal structures including compartment, TAD and loop. Following 1,6-HD treatment, compartment changes were subdivided into four compartment change-types based on the changes of the PC1 value: strengthened, stable, weakened, and flipped compartment (Figure 5A). As shown in Figure 5B and S5A, the B compartment is more stable after 1,6-HD treatment compared to the A compartment. The changes of the inter-compartment interaction also showed that compartment A is more dynamic than B (Figure 5C and 5D). We found that the number of interactions between A-A compartments, even at distances as long as 100Mb, significantly increased. In contrast, although the number of short distance B-B interactions also increased, long distance B-B interactions were stable (Figure 5C and 5D).

**Figure 5.**
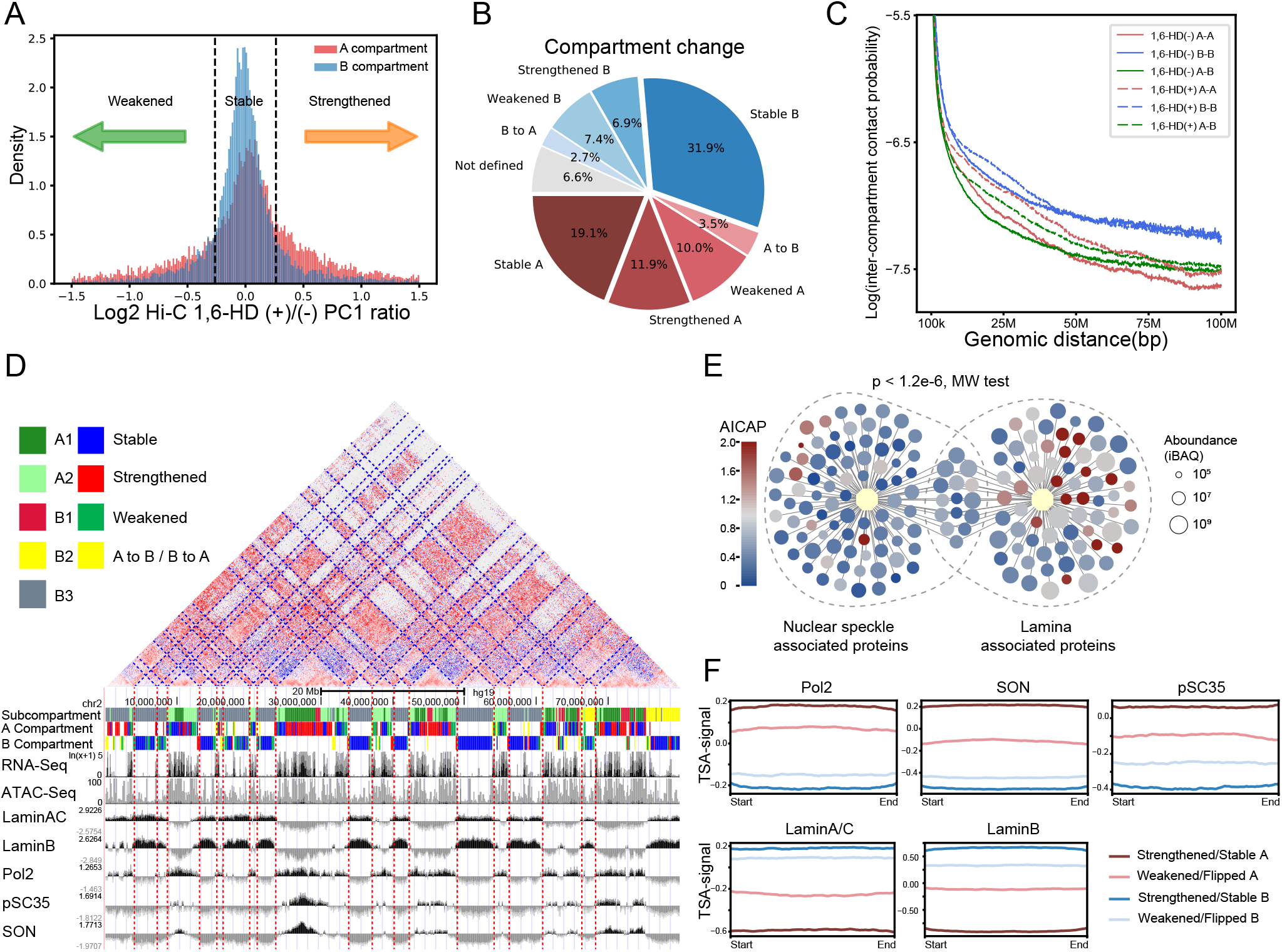
Chromatin compartment changes are related to neighboring nuclear condensates. **A.** A(red) and B(blue) compartments can be classified into 4 compartment change-types as strengthened, stable, weakened based on their PC1 value ratio (1,6-HD+/−). 20% was chosen as the threshold for distinguishing stability or not. **B.** The fraction of four kinds of compartment change-types. **C.** Contact probability between compartments along genomic distance. **D.** Examples of strengthened/stable compartments and corresponding nuclear speckle/lamina TSA-seq plots. The plotting of log2 ratio of TSA read density versus input read density was used to measure the distance from a chromatin region to a specific nuclear condensate. Chr2: 0-80M. **E.** AICAP values of nuclear speckle and lamina associated proteins. Data resource (Cutler et al., 2019; Dopie et al., 2020). **F.** Aggregation analysis of TSA-seq signal enrichment of compartment change-types.

To identify the causal factors responsible for the observed compartment changes, we analyzed the relationship between the compartment change-types and their neighbor nuclear condensates. Nuclear speckle is a typical nuclear condensate characterized by active gene expression and RNA splicing. In comparison, nuclear lamina constructs a stable framework that attach heterochromatin[31]. SON and pSC35 are two hallmarks of nuclear speckles[32, 33], while lamin B and A/C are two hallmarks of nuclear lamina[34]. Previous studies have used proximate labeling to capture proteins located proximally to the nuclear speckle and lamina, and obtained high confidence subsets by comparing appropriate control samples[35, 36]. We displayed AICAP values and protein contents for these high confidence subsets proteins to show the LLPS properties of these nuclear condensates. As shown in Figure 5E, proteins included in nuclear speckles exhibit low AICAP values, characterized by a median value of 0.50. In comparison, proteins included in nuclear lamina possess high AICAP values, with a median value of 0.81.

TSA-seq was a recently developed technique for estimating the average chromosomal distances from nuclear speckles to nuclear lamina[37]. Using TSA-seq data, we plotted the distance of each compartment change-type to nuclear speckle and lamina. As shown in Figure 5D, following 1,6-HD treatment, the compartments located closer to the nuclear lamina were generally stable, while the compartments located closer to nuclear speckles tend to be enhanced. Aggregation analysis showed that strengthened/stable A, flipped A, weakened/flipped B and strengthened/stable B sequentially distributed between nuclear speckles and nuclear lamina (Figure 5F).

Compartment changes were even constrained by neighbor compartment and nuclear condensate. Generally, compartment A locate coincide with nuclear speckles while compartment B locate coincide with nuclear lamina. We found that for the strengthened and stable compartments, their neighbors were often the same types of compartments, while for the weakened and flipped compartments, their neighbors were often different types of compartments (Additional file 1: Figure S5B). Meanwhile, following 1,6-HD treatment, the compartments undergoing a conversion from A to B were usually surrounded by B compartments, while the compartments undergoing B to A conversion were commonly surrounded by A compartments (Additional file 1: Figure S5B and S5C).

Together, these results indicate that compartment A are more sensitive to 1,6-HD treatment, and this sensitivity were in agree with the AICAP of proximate nuclear condensates as well. This result is also consistent with one recent study showing that heterochromatic regions exhibit stronger internal attractions than euchromatin[38].

### Nuclear condensates affect various levels of chromatin organization

To further evaluate the role of LLPS in finer chromatin structure, we next investigated the chromatin organization changes at TAD and loops level following 1,6-HD treatment. The inter-TAD interaction increased significantly at a number of sites (example ‘a, b’ in Figure 6A), which resulted in the loss of 15.6% of the TAD boundaries (Additional file 1: Figure S6A). Compared with the lost boundaries, CTCF and cohesin peaks were significantly enriched at the stable boundaries (Figure 6B), and tandem strong CTCF peaks coexist in the same boundary (example ‘c’ in Figure 6A). Furthermore, the sensitivity of TAD to 1,6-HD treatment showed a sub-comp-dependent manner. As shown in Figure 6C, the order of intra and inter-TAD interactions increment was A1>A2B1>B2B3. Next, we combined TSA-seq to investigate the distance of TAD boundaries to nuclear speckle and lamina. As shown in Figure 6D, the stable TAD boundaries localized closer to nuclear lamina, while the lost TAD boundaries localized closer to Pol2 and nuclear speckle. This distribution indicated that chromatin TAD stability was also related to the LLPS properties of neighbor nuclear condensates.

**Figure 6.**
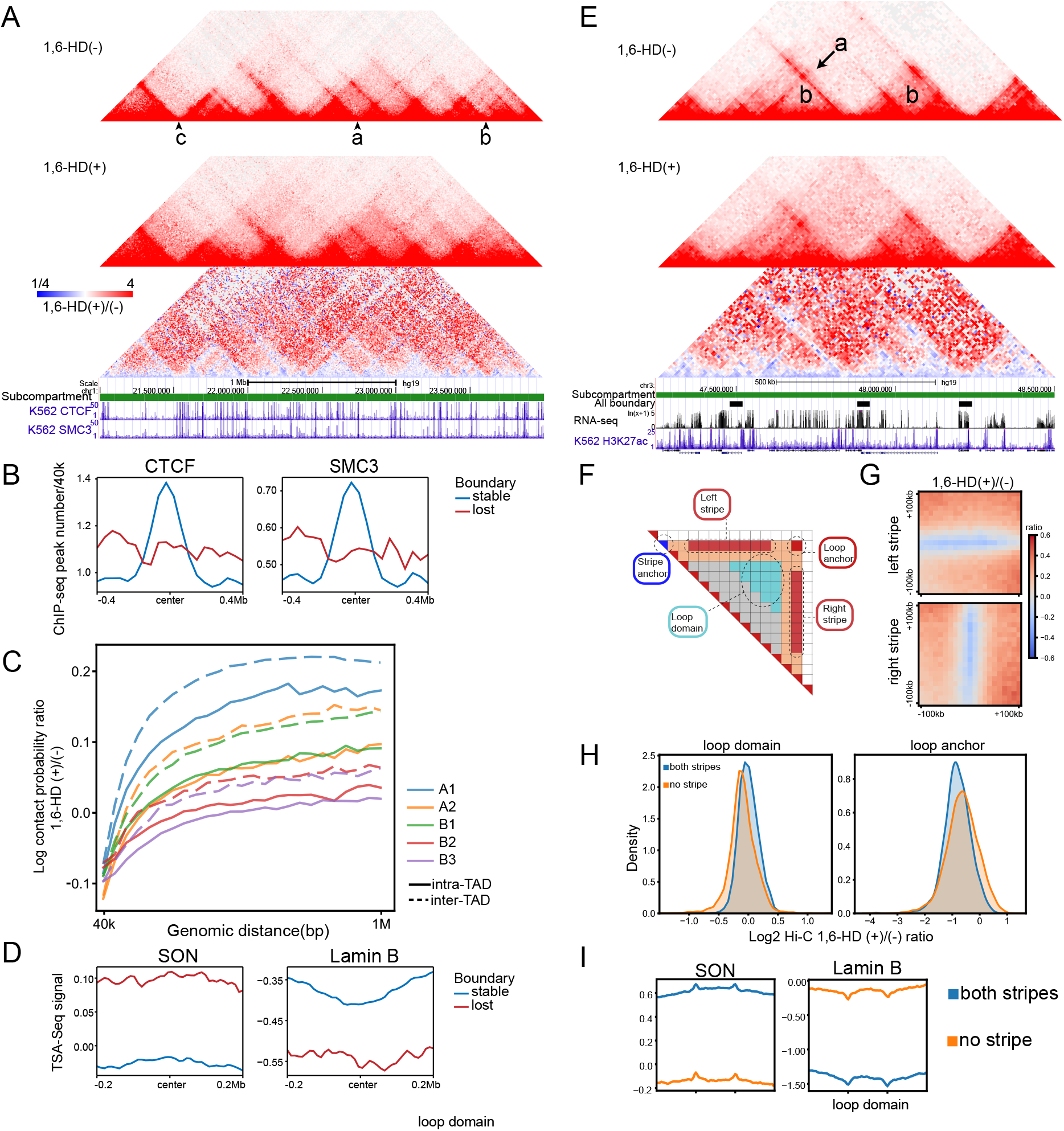
Chromatin TAD and loop changes are related to neighboring nuclear condensates. **A.** Examples of stable/lost TAD boundaries. example ‘a, b’ lost, example ‘c’ stable. **B.** Aggregation analysis of CTCF/SMC3 ChIP-seq peaks at stable/lost TAD boundaries. **C.** Intra−/inter-TAD interaction changes (1,6-HD(+)/(−) ratio) in different sub-compartments. Interaction between different sub-compartments were skipped. **D.** Aggregation analysis of TSA-seq signal at stable/lost TAD boundaries. **E.** Examples of loop anchor interaction, loop domain and DNA stripe after 1,6-HD treatment. **F.** Schematic illustration of loop anchor interaction, loop domain, DNA stripe and stripe anchor. **G.** Left and right stripe signal aggregation at 10kb resolution. **H.** Aggregation analysis of loop domain (left) and loop anchor interaction(right) signal changes after 1,6-HD treatment. “both/no stripe” indicates both or no overlap between loops anchor and stripe anchor. **I.** Aggregation analysis of TSA-seq signal at loop with “both/no

1,6-HD treatment caused drastic changes on DNA loops. The overall strength of the loops decreased 60% on average (Additional file 1: Figure S6B), and 61.5% of all DNA loops disappeared after 1,6-HD treatment (Additional file 1: Figure S6C), which is much more than that of TAD boundary. DNA stripe is a subunit of DNA loop which is considered to be the prototypes of cohesin extrusion, and nearly 80% of stripe domains were associated with active enhancers in mouse B cells[39]. After 1,6-HD treatment, both left and right DNA stripes were visibly weakened (example ‘a’ in Figure 6F, Figure 6G), and 40% of the DNA stripes disappeared (Additional file 1: Figure S6C). However, interactions surrounding the DNA stripes (loop domain, Figure 6F and example ‘b’ in Figure 6E) appeared to be increased (Figure 6G). Since stripe anchors often locate coincide with loop anchors, DNA loops can be divided into two types, which are loops carrying two or none DNA stripes. We therefore tested the stability of these two types of DNA loops. As shown in Figure 6H, DNA loops containing DNA stripes (both stripes) weakened more than the loops not containing DNA stripes (no stripe). When analyzed together with the TSA-seq data, we further found that the loops containing DNA stripes localized closer to nuclear speckles and farther to lamina compared with loops not associated with DNA stripes (Figure 6I).

In summary, these results strongly indicated that chromatin changes in TADs and loops were related to proximate nuclear condensates. Taken the compartment changes into account, we revealed that the 3D chromatin organization in the transcriptionally active region are more sensitive to 1,6-HD treatment at various levels.

## Discussion

To quantifying LLPS-dependence of proteins and 3D genome structure, we disrupted hydrophobicity dependent LLPS properties by 1,6-HD treatment, then developed Hi-MS and performed Hi-C to quantify changes in both chromatin binding proteins and higher-order chromatin structure in K562 cell. We proved that AICAP is a powerful and reliable tool for measurement of LLPS property.

### LLPS is an intrinsic physicochemical property of active biological processes

When we compared the AICAP values of different proteins in the same cell type, we found that the dynamic regulatory components, both protein (trans) and DNA (cis), exhibited low AICAP values, which demonstrates that physicochemical properties and biological functions are well correlated. Our data provided vivid examples for the views of Banani et al., who claimed that “one important advantage of phase-separated structures is that all these potential functions can be switched on and off extremely rapidly by controlling the formation and dissolution of a condensed phase”[40]. The consequences of losing this structure and function coordination are likely to be catastrophic for cells. If the constitutive and structural proteins (compartment B) exhibit a low AICAP value (more LLPS property), they will be unstable when exposed to external interference. Otherwise, if the regulatory proteins (compartment A) exhibit a high AICAP value (less LLPS property), they will only respond to strong stimuli, and cannot respond sensitively to external changes.

It was previously reported that heterochromatin binding protein HP1a undergo phase-separation[8]. However, we show here that HP1a exhibits a high AICAP value. Fortunately, Strom et al. also reported that heterochromatin may be initially formed via LLPS, but it gradually matures into an immobile structure no longer sensitive to 1,6-HD treatment[8]. This liquid-solid transition may be necessary for heterochromatin to inhibit transposon activity and maintain the structural stability of the genome. This transition further supported our conclusion that active biological processes exhibit more LLPS property.

### AICAP values is directly proportional to proteins with distinct functions in the same category

As shown above, the change in AICAP values is directly proportional to the transition from active to repressed chromatin states. Unexpectedly, the AICAP value of each member in the same functional category is not identical, with some proteins varying considerably (Figure 4B). This variance might reflect differences in function of these proteins. For instance, while histone family exhibit AICAP values of approx. 1.0, the H1 and H2B family exhibit relatively low AICAPs. H1 was recently reported to enhance nuclear condensates via LLPS [41], [42], and H2B ubiquitylation was reported to decondense chromatin and facilitate ‘crosstalk’ between H2B and subsequent histone modification and variants[43]. The accessibility of H1 and H2B may agree with their relatively low AICAPs.

The structural proteins ZNF143 exhibit an AICAP value of only 0.17, far below other chromatin structural proteins. Although ZNF143 was reported to be a structural protein, it preferentially localizes in active transcriptional regions[44]. The low AICAP may facilitate the cooperation of ZNF143 with other transcription-related proteins. Furthermore, BRD4 exhibit a much higher AICAP value than MED1. Functionally, BRD4 bind SEs earlier than MED1 to drive out HP1a and decondensed local DNA, this function requires strong DNA binding capacity and stability; however, MED1 binds open SEs, connects TF and RNAPII through LLPS, and initiation transcription[45]. We also showed that GATA1 exhibits a higher AICAP than GATA2. Functionally, a high AICAP value indicates high DNA binding affinity. This property may be required by ‘GATA switch’ process during hematopoietic cell development, where GATA1 replaces GATA2 on the GATA motifs[46].

We show that heterochromatin-related proteins usually exhibit high AICAP values, a fact that consistent with the structural function and stability of heterochromatin. However, histone lysine methyltransferases (GLP/G9A), which control PRC2 recruitment and H3K27 trimethylation[47], exhibited AICAP values proximate 0.2. Similarly, the de novo methylases DNMT3B have lower indexes than DNMT1 which preserve methylation, even though they are both DNA methylases. De novo methylation is a dynamic process, while methylation maintaining is relatively stable process[48]. All of these AICAP-function relationships support the LLPS nature of active/regulatory biological processes.

### Limitations and suggestions for future research

Although we have obtained AICAP values for thousands of proteins and proved that they are closely related to the chromatin organization stability, histone modification may also play a critical role in the local hydrophobicity-dependent chromatin condensation[42, 49]. Histone acetylation disrupts chromatin droplets and re-phase-separate by multi-bromodomain proteins, including the transcriptional regulator BRD4[42]. Recent evidence also suggest that histone modifications affect local hydrophobic interactions[50]. Histone variants also play roles in fine-tuning chromatin organization and function[51]. However, because histones are tightly bound to DNA, it is difficult for Hi-MS to determine how different modifications change the hydrophobic interactions within a local DNA environment.

Proteins with an AICAP value larger than 1.0 are generally bound to chromosomes non-physiologically. However, a small number of these proteins may enter the cell nucleus during specific cell cycle or state. For example, cyclin B (CCNB1, AICAP value=1.6) enter the nucleus during the G2/M transition and STAT1(AICAP value=8.01) enter the nucleus following interferon stimulation. Since the active transport of these proteins requires passing through NPC, while NPC channel has been reported to be a hydrophobic environment, these proteins may also possess strong hydrophobic-dependent LLPS potential. However, these proteins need to be studied under specific conditions, in order to effectively distinguish these proteins from non-physiologically binding proteins.

In this study, we enriched regulatory genome associated proteins using our newly developed Hi-MS method. Future studies should target marker proteins directly using specific antibody, thus target specific condensate in the cytoplasm or nucleus, and find key factors for driving the separation in each type of condensate.

## Methods

### Cells and Cell Culture

Human female K562 cells were obtained from ATCC(Cat#CCL-243) and cultured in RPMI 1640 medium containing 10% FBS and 1% penicillin/streptomycin. All cultures were incubated at 37°C in 5% CO2.

#### 1,6-Hexanediol treatment

For isotonic condition, 1,6-Hexanediol (Sigma Cat#240117) was dissolved in RPMI 1640 medium containing 10% FBS to a concentration of 10% to make a storage solution. The working solution was made by dilution using RPMI 1640 medium containing 10% FBS immediately before use. For hypotonic condition, 1,6-Hexanediol was dissolved in H2O to a concentration of 30% to make a storage solution. The working solution was made by 1:2 mix of 30% storage solution and RPMI 1640 medium containing 10% FBS immediately before use.

### Hi-C

The BL-Hi-C library construction was performed as previously described with some modifications[22]. 10^6^ K562 cells were incubated with 1% formaldehyde in PBS to crosslink protein–DNA in the cells, then, the cells were suspended using lysis buffer (50 mM HEPES-KOH, 150mM NaCl, 1 mM EDTA, 1% Triton X-100, and 0.1% SDS). The genome was then digested by HaeIII (NEB) into fragments with blunt-ends. The DNA fragments were treated with adenine and ligated with bridge linker with biotin for 4h at RT. The unligated DNA fragments were digested with exonuclease (NEB). Next, the cells were digested by Proteinase K (Ambion) overnight, and the DNA was purified using phenol–chloroform (Solarbio) extraction with ethanol precipitation. Then, the ligated DNA was fragmented into 300bp using an S220 Focused-ultrasonicator (Covaris), and the biotin-labeled DNA fragments were enriched by Dynabeads M280 beads (Thermo Fisher). The enriched DNA library was amplificated by PCR using Q5 DNA polymerase (NEB). After size-selection with the AMPure XP beads (Beckman, Germany), the libraries were sequenced on an Illumina HiSeq 2500 sequencer.

#### Data processing

Bridge linkers were trimmed using ChIA-PET2 software [52] using parameters “-A ACGCGATATCTTATC -B AGTCAGATAAGATAT -k 2 -m 1 -e 1”. The resulting clean paired-end reads were aligned independently to hg19 human genome using bwa mem and then processed by HiC-Pro software [53] to obtain valid interaction pairs (“.validPairs”) and subsequent matrix of different resolution. “.hic” files of two condition were converted from “.validPairs” using Juicer Tools 1.13.02 and used for subsequent analysis.

#### Compartment calling

A/B compartment were identified by eigenvector decomposition on the Pearson’s correlation matrix of KR-balanced OE (observed/expected) cis-interaction matrix at 100kb resolution. The positive and negative values of first eigenvector (PC1) for each 100kb bin were assigned to A(active) and B(inactive) compartments based on its association with gene density. PC1 ratio = (Hex+ PC1 value) / (Hex− PC1 value). The positive value represented that the PC1 value increased after 1,6-hexanediol treatment, suggesting the A/B compartment feature became strengthened. In opposite, the negative value indicated the A/B compartment feature became weakened. We took the change within ±20% as stable A/B compartment and those beyond ±20% as weakened/strengthened. Compartments with different signs before and after treatment were annotated as flipped compartments.

#### Saddle plot

To measure the strength of compartments, interaction bins of KR-balanced OE cis-interaction matrix at 40kb resolution were sorted according to PC1 value. All cis interactions with similar PC1 values were aggregated to obtain compartmentalization saddle plot with preferential B-B interactions in the upper left corner and preferential A-A interactions in the lower right corner.

#### Subcompartment annotation

Rao et al. divided the A/B compartment into five subcompartments namely A1, A2, B1, B2, B3 based on the regions’ inter-chromosome Hi-C interaction in GM12878 cells, which required as high as 1kb resolution. Different genomic and epigenetic features were observed in different subcompartments. Xiong et al. developed SNIPER to accurately infer subcompartments based on Hi-C data of moderate depth (~500 million mapped reads). Here we utilized the SNIPER annotation of subcompartment in K562 cells[54].

#### Topological associated domain boundary

TAD boundaries were identified using KR-balanced matrix at 40 kb resolution by a Perl script matrix2insulation.pl (https://github.com/dekkerlab/crane-nature-2015) as previously described[55]. The insulation scores were calculated for each chromosome bin by a sliding 1Mb x 1Mb square along the diagonal of the matrix. A 200kb window was used for calculation of the delta vector. TAD boundaries with “Boundary strength” under 0.1 were filtered. TAD boundaries whose centers located within ±80kb (2 bins) in two conditions were defined as unchanged boundary.

#### Loop calling

Loops were called using HICCUPS [56] at 5/10/20kb resolution with parameters “-k KR -f 0.1,0.1,0.1 -p 4,2,1 -i 7,5,3 -t 0.02,1.5,1.75,2 -d 20000,20000,50000”. Loops detection before and after treatment were conducted separately and differential loops were annotated as loops that were not detected after treatment using bedtools pairToPair (loops anchor were slopped with 10kb to avoid false positive).

Aggregation peak analysis (APA) were generated at 5kb using Juicer APA subcommand with slight modification. Loops were grouped based their subcompartments and the resulted APA matrix were divided by their corresponding number of loops.

Loop signal change were defined as normalized Hi-C contact probability ratio at loop pixels.

#### Stripe calling

Stripes were identified using the R script provided by Aleksandra et al. as previous described [39].The analyses were performed using raw interaction matrices and the normalized matrices generated using juicer software (the .hic files). The matrices were exported to a .txt format from the .hic files using the dump function of juicer. The stripe calling was implemented and performed in R using custom functions.

### Hi-MS (chromatin associated protein capture)

The Hi-MS sample was prepared based on BL-Hi-C protocol to extract chromatin associated proteins. 107 K562 cells were incubated with 1% formaldehyde in PBS to crosslink protein–DNA in the cells, then, the cells were suspended using 1% SDS lysis buffer (50 mM HEPES-KOH, 150mM NaCl, 1 mM EDTA, 1% Triton X-100, and 1% SDS). After wash cells with cutsmart buffer with 1%TX-100, the genome was then digested by HaeIII (NEB) into fragments with blunt-ends. The DNA fragments were treated with adenine and ligated with bridge linker with biotin for 4h at RT. Then, the cells were washed by 0.2%SDS nucleus lysis buffer (20mM Tris-HCl, 50mM NaCl, 2mM EDTA, 0.2% SDS, 1×protease inhibitor) once, then incubate in 0.2%SDS nucleus lysis buffer at 4°C overnight. The next morning, the cells were washed once again and resuspended in 0.2%SDS nucleus lysis buffer. Cells were sonicated using Digital Sonifier Cell Disruptor at 40% output for 24 cycles, each 5s ON and 5s OFF. After sonication, 2x volumes of IP dilution buffer (20mM Tris pH8, 2mM EDTA, 450mM NaCl, 2% Triton X-100, protease inhibitors) was added and incubate for 1hrs at 4C with rotation. The biotinylated linker/DNA/protein complex in supernatant was then incubated with 1ml M280 magnet beads slurry (Thermo-Fisher Cat#60210) for 2hrs at 4°C with rotation. Beads were then washed 3 times with cold IP wash buffer 1 (20mM Tris pH8, 2mM EDTA, 50mM NaCl, 1% Triton X-100, 0.1% SDS), once with cold TE buffer (1mM Tris pH8, 1mM EDTA). The complex were eluted twice for 5min at 100 °C in 60ul H2O each time and sent for label-free quantitative mass spectrometry (MS) analysis.

### Hi-MS (Protein Sample Preparation for Mass Spec analysis)

#### In-gel digestion of proteins (condition 1)

The gel bands containing the protein sample were manually excised. Each of the protein bands was then digested individually as below. The protein bands were cut into small plugs, washed twice in 200 μl of distilled water for 10 min each time. The gel bands were dehydrated in 100% acetonitrile for 10 min and dried in a Speedvac for approximately 15 min. Reduction (10 mM DTT in 25 mM NH4HCO3 for 45 min at 56°C) and alkylation (40 mM iodoacetamide in 25 mM NH4HCO3 for 45 min at room temperature in the dark) were performed, followed by washing of the gel plugs with 50% acetonitrile in 25 mM ammonium bicarbonate twice. The gel plugs were then dried using a speedvac and digested with sequence-grade modified trypsin (40 ng for each band) in 25 mM NH4HCO3 overnight at 37 °C. The enzymatic reaction was stopped by adding formic acid to a 1% final concentration. The solution was then transferred to a sample vial for LC-MS/MS analysis.

#### LC-MS/MS analysis (condition 1)

All nano LC-MS/MS experiments were performed on a Q Exactive (Thermo Scientific) equipped with an Easy n-LC 1000 HPLC system (Thermo Scientific). The peptides were loaded onto a 100 μm id×2 cm fused silica trap column packed in-house with reversed phase silica (Reprosil-Pur C18 AQ, 5 μm, Dr. Maisch GmbH) and then separated on an a 75 μm id×20 cm C18 column packed with reversed phase silica (Reprosil-Pur C18 AQ, 3 μm, Dr. Maisch GmbH). The peptides bounded on the column were eluted with a 78-min linear gradient. The solvent A consisted of 0.1% formic acid in water solution and the solvent B consisted of 0.1% formic acid in acetonitrile solution. The segmented gradient was 4–8% B, 8 min; 8–22% B, 50 min; 22–32% B, 12 min; 32-90% B, 1 min; 90% B, 7min at a flow rate of 300 nl/min.

The MS analysis was performed with Q Exactive mass spectrometer (Thermo Scientific). With the data-dependent acquisition mode, the MS data were acquired at a high resolution 70,000 (m/z 200) across the mass range of 300–1600 m/z. The target value was 3.00E+06 with a maximum injection time of 60 ms. The top 20 precursor ions were selected from each MS full scan with isolation width of 2 m/z for fragmentation in the HCD collision cell with normalized collision energy of 27%. Subsequently, MS/MS spectra were acquired at resolution 17,500 at m/z 200. The target value was 5.00E+04 with a maximum injection time of 80 ms. The dynamic exclusion time was 40s. For nano electrospray ion source setting, the spray voltage was 2.0 kV; the heated capillary temperature was 320 °C.

#### In-solution digestion of proteins (condition 2)

Protein concentration was determined by Bradford protein assay. Extracts from each sample (40◻μg protein) was reduced with 10◻mM dithiothreitol at 56◻°C for 30◻min and alkylated with 10◻mM iodoacetamide at room temperature in the dark for additional 30◻min. Samples were then digested using the filter-aidedsample preparation (FASP) method with trypsin[57]; tryptic peptides were separated in a home-made reverse-phase C18 column in a pipet tip. Peptides were eluted and separated into nine fractions using a stepwise gradient of increasing acetonitrile (6%, 9%, 12%, 15%, 18%, 21%, 25%, 30%, and 35%) at pH 10. The nine fractions were combined to six fractions, dried in a vacuum concentrator (Thermo Scientific), and then analyzed by liquid chromatography tandem mass spectrometry (LC-MS/MS).

#### LC-MS/MS analysis (condition 2)

Samples were analyzed on Orbitrap Fusion Lumos Plus mass spectrometers (Thermo Fisher Scientific, Rockford, IL, USA) coupled with an Easy-nLC 1000 nanoflow LC system (Thermo Fisher Scientific). Dried peptide samples were re-dissolved in Solvent A (0.1% formic acid in water) and loaded to a trap column (100◻μm◻×◻2◻cm, home-made; particle size, 3◻μm; pore size, 120◻Å; SunChrom, USA) with a max pressure of 280◻bar using Solvent A, then separated on a home-made 150◻μm◻×◻12◻cm silica microcolumn (particle size, 1.9◻μm; pore size, 120◻Å; SunChrom, USA) with a gradient of 5–35% mobile phase B (acetonitrile and 0.1% formic acid) at a flow rate of 600◻nl/min for 75◻min. For detection with Fusion Lumos mass spectrometry, a precursor scan was carried out in the Orbitrap by scanning m/z 300−1400 with a resolution of 120,000 at 200◻m/z. The most intense ions selected under top-speed mode were isolated in Quadrupole with a 1.6◻m/z window and fragmented by higher energy collisional dissociation (HCD) with normalized collision energy of 35%, then measured in the linear ion trap using the rapid ion trap scan rate. Automatic gain control targets were 5◻×◻105◻ions with a max injection time of 50◻ms for full scans and 5◻×◻103 with 35◻ms for MS/MS scans. Dynamic exclusion time was set as 18◻s. Data were acquired using the Xcalibur software (Thermo Scientific).

The mass spectrometry proteomics data have been deposited to the ProteomeXchange Consortium via the PRIDE partner repository [58] with the dataset identifier PXD021434. Data processing and protein quantification. All the MS data were processed in the Firmiana database (Feng et al., 2017). Raw files were searched against the human National Center for Biotechnology Information (NCBI) Refseq protein database (updated on 07-04-2013, 32015 entries) by Mascot 2.3 (Matrix Science Inc). The mass tolerances were 20 ppm for precursor and 0.05 Da or 0.5 Da for productions for Q Exactive (Experiment 1) and Fusion (Experiment 2) respectively. Up to two missed cleavages were allowed. The data were also searched against a decoy database so that peptide identifications were accepted at a false discovery rate (FDR) of 1%. Proteins with at least 1 unique peptide with Mascot ion score greater than 20 or 2 peptides with Mascot ion score greater than 20 were remained. Label-free protein quantifications were calculated using a label-free, intensity based absolute quantification (iBAQ) approach (Schwanhausser et al., 2011). The fraction of total (FOT) was used to represent the normalized abundance of a particular protein/peptide across control and treated samples. FOT of protein was defined as a protein’s iBAQ divided by the total iBAQ of all identified proteins within one sample. The FOT was multiplied by 106 for the ease of presentation. The missing data were imputed with the minimum values. After missing value imputation, quantile normalization was applied.

#### Statistical analysis

iBAQ values were used in the comparison between control and mock samples and FOT were used in the comparison between control and treated samples. P value was calculated to measure the statistical significance of protein abundance difference of each identified protein in the replicate experiments by t test. (Additional file 2: Table S1)

### Immunofluorescence

Coverslips were coated at RT with 5ug/mL Poly-L-lysine solution (Sigma-Aldrich, P4707) for 30 minutes. K562 Cells were plated on the pre-coated coverslips and grown for 1 hr followed by fixation using 4% paraformaldehyde (Sigma Aldrich, 47608) in PBS for 10 minutes. Then the cells were permeabilized using 0.5% Triton X-100 (Sigma Aldrich, X100) in PBS for 10 minutes. Cells were blocked with 4% Bovine Serum Albumin, BSA, (VWR, 102643-516) for 1h and the indicated primary antibody was added at suitable concentration in PBST for 1 hour. Cells were washed with PBS three times followed by incubation with secondary antibody at a concentration of 1:1000 in PBS for 1 hour. After washing twice with PBS, Cells were washed once in water followed by mounting the coverslip onto glass slides with Vectashield (VWR, 101098-042) and finally sealing the coverslip with nail polish (Electron Microscopy Science Nm, 72180). Images were acquired at an Zeiss LSM 880 confocal microscope with 63×objective using ZEN acquisition software. Images were post-processed using Fiji Is Just ImageJ (FIJI). Antibodies used were listed in Additional file 6: Table S5.

### RNA-seq

Total RNA was extracted from the K562 cells using TRIZOL (Ambion, USA). The library construction and sequencing were performed by ANOROAD (China). Reads were aligned to hg19 genome using hisat2 and resulted sam files were sorted using samtools. The expression profiles were generated using cufflinks cuffnorm with geometric normalization. Signal tracks were produced by deeptools bam Coverage command.

### ATAC-seq

The ATAC library was prepared using Omni-ATAC protocol as previously described. Briefly, 50,000 cells were Pellet and resuspend using 50 ul cold ATAC-Resuspension Buffer (RSB) (10 mM Tris-HCl, 10 mM NaCl, 3 mM MgCl_2_, pH 7.4) containing 0.1% NP40, 0.1% Tween-20, and pipette up and down 3 times. Incubate on ice for 3 minutes. Wash out lysis with 1 ml of cold ATAC-RSB containing 0.1% Tween-20 but NO NP40 and invert tube 3 times to mix. Pellet and Resuspend cells in 50 ul of transposition mixture by pipetting up and down 6 times. The nuclei were then incubated with the Tn5 Transposition mix (10 ul 5x TTBL buffer, 3 ul TTE Mix V50 transposase, 16.5 ul PBS, 0.5 ul 10% Tween-20, 20 ul H2O) at 37◻°C for 30◻min (TruePrep® DNA Library Prep Kit V2 for Illumina, Vazyme, China). After the tagmentation, the stop buffer was directly added to the reaction to end the tagmentation. PCR was performed to amplify the library in 12 cycles. After the PCR reaction, the libraries were purified with 1.2× AMPure beads (Beckman, Germany). The libraries were sequenced using an Illumina HiSeq 2500 sequencer.

ATAC-seq raw reads were trimmed to remove adaptor sequence and mapped to hg19 genome with Bowtie2 using parameters “--very-sensitive -X 2000” and duplicates were removed using Picard Mark Duplicates command. Signal tracks were produced by deeptools bam Coverage command. Peaks were called by MACS2 using parameters “--nomodel --shift -100 --extsize 200 -B --call-summits –SPMR”.

### Restriction endonuclease recognition motif frequency

Recognition motif GGCC (HaeIII) and GATC (DpnII and HindIII) frequency were defined as occurrence times per 500 bp across genome, which was transform to bigwig coverage by deeptools bam Coverage. The signal at H3k27ac/EZH2 ChIP-seq peaks were generated using deeptools compute Matrix.

## Acknowledgments

We thank Wei Xie, Pi-long Li, and Hai-tao Li at Tsinghua U. for useful discussion and critical reading of the manuscript.

## Authors’ contributions

Conceptualization, M.L.S., T.T.L and Y.C.; Methodology, M.L.S., T.T.L, Y.C. and Z.Y.L; Investigation, M.L.S., K.Q.Y., C.H, T.Y.C., M.W.L, and J.F.W.; Formal Analysis, K.Q.Y.; Data Curation, K.Q.Y., Visualization, K.Q.Y., Writing – Original Draft, M.L.S. T.T.L. and K.Q.Y.; Writing – Review & Editing, M.Q.Z.; Funding Acquisition, M.Q.Z. T.T.L. and Y.C.; Resources, M.Q.Z., T.T.L., Y.C., T.T.W., and J.Q.,; Supervision, M.Q.Z. and T.T.L.

## Ethics approval and consent to participate

Not applicable.

## Consent for publication

Not applicable.

## Figure titles and legends

**Figure S1.**
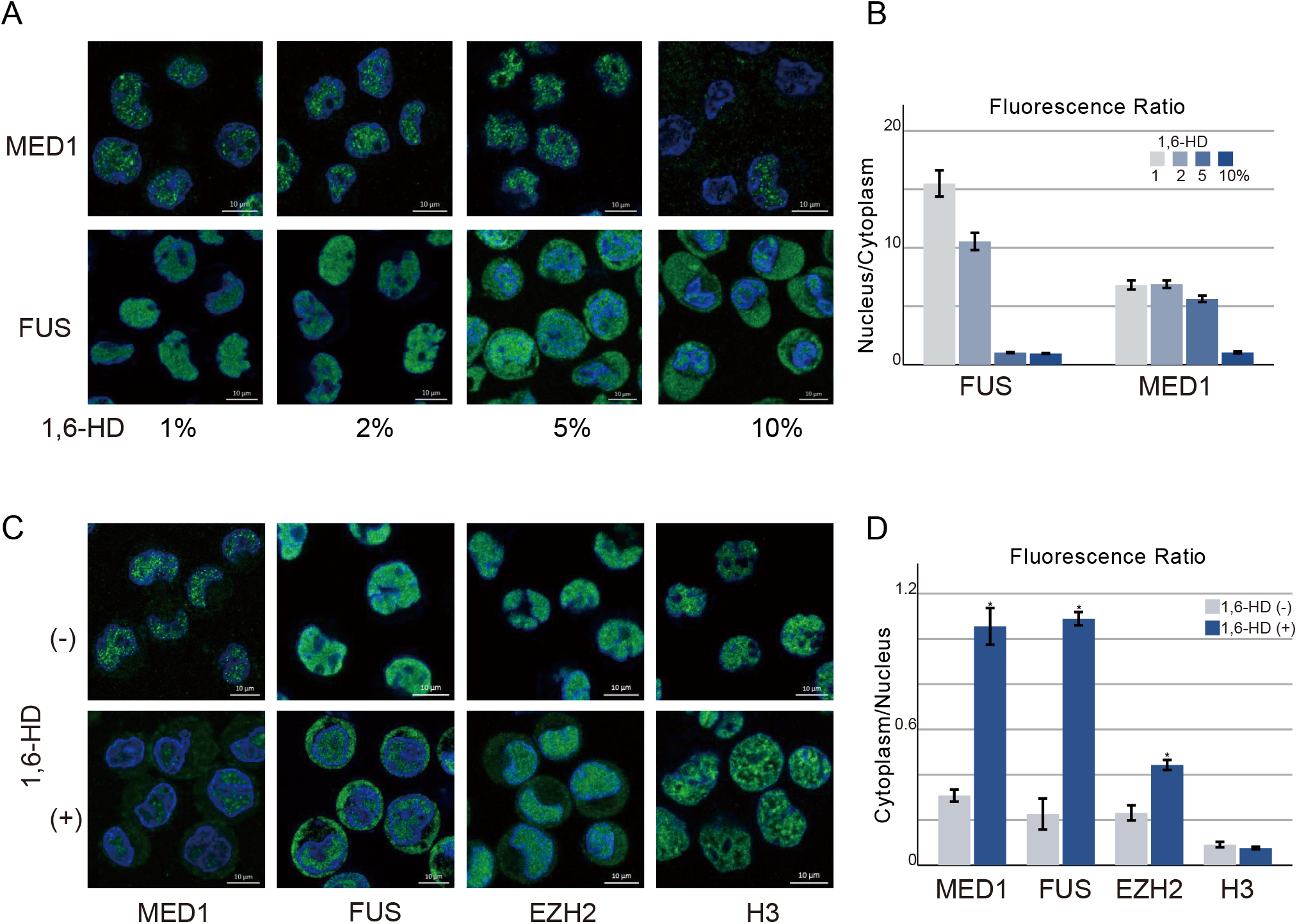
Proteins exhibit different sensitivities to 1,6-HD treatment. **A.** MED1/FUS distribution at 1,6-HD concentration from 1-10% in K562 cells. Incubation time: 20min. Blue: DAPI, green: MED1/FUS. Scale bar: 10μm. **B.** Quantification of nuclear/cytoplasmic fluorescence signal for MED1/FUS upon 1,6-HD treatment (n = 100), as in (A). **C.** Nuclear-cytoplasmic distribution of MED1, FUS, EZH2 and H3K4Me3 after 10% 1,6-HD treatment for 20 mins in hypotonic condition. **D.** Quantification of cytoplasmic/nuclear fluorescence signal for MED1, FUS, EZH2 and H3 upon 1,6-HD treatment (n = 100), as in (B).

**Figure S2.**
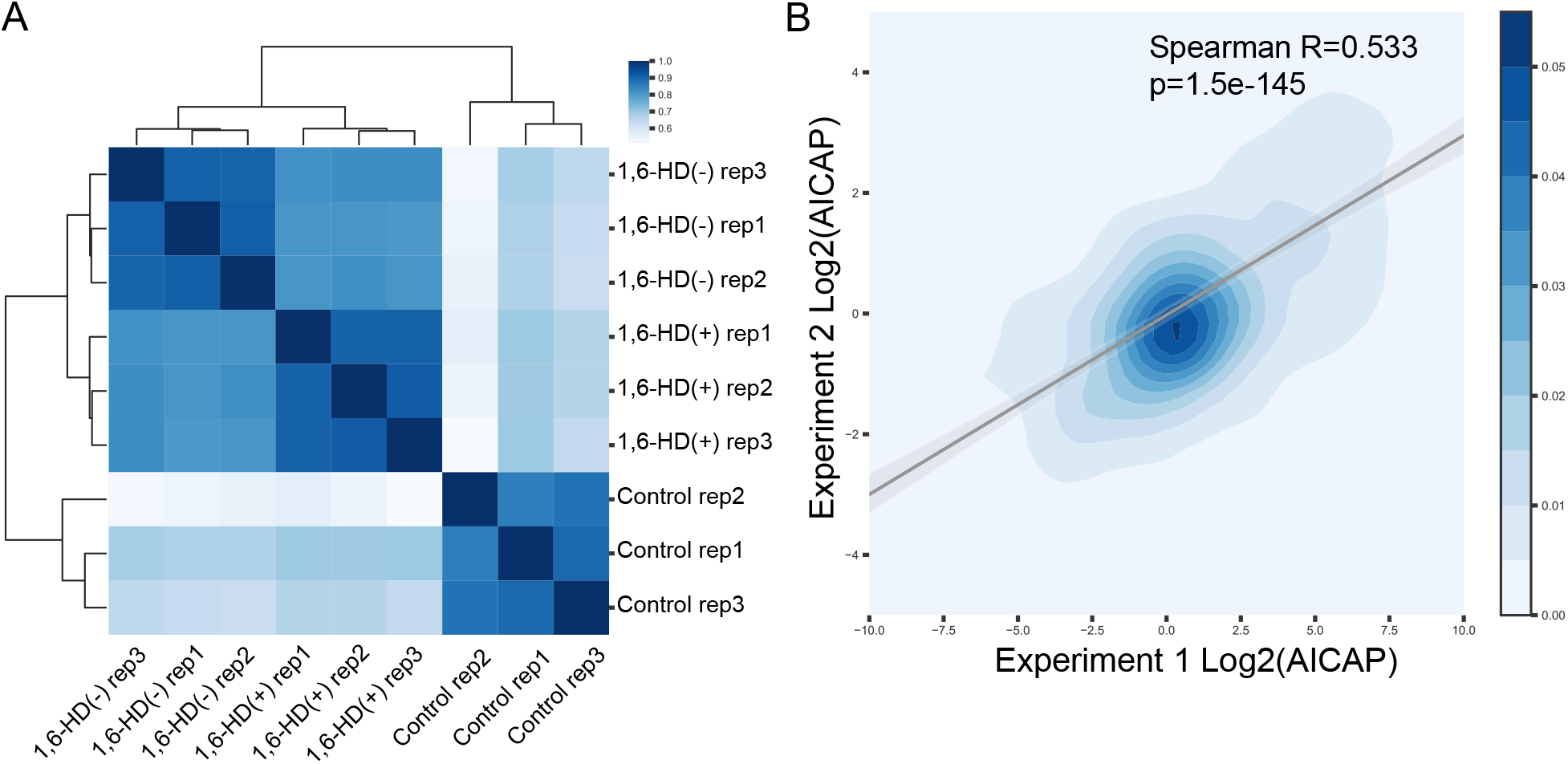
Reproducibility of Hi-MS and AICAP. **A.** Clustering of replicated Hi-MS samples using Spearman correlation of protein content (iBAQ signal). (−): wild type samples, (+): 1,6-HD treated samples and control: undigested control samples. **B.** Spearman correlation of AICAP values of two independent experiments. They were prepared from different batches of cells, different types of digestion (in gel or in solution) and different types of mass spectrometers.

**Figure S3.**
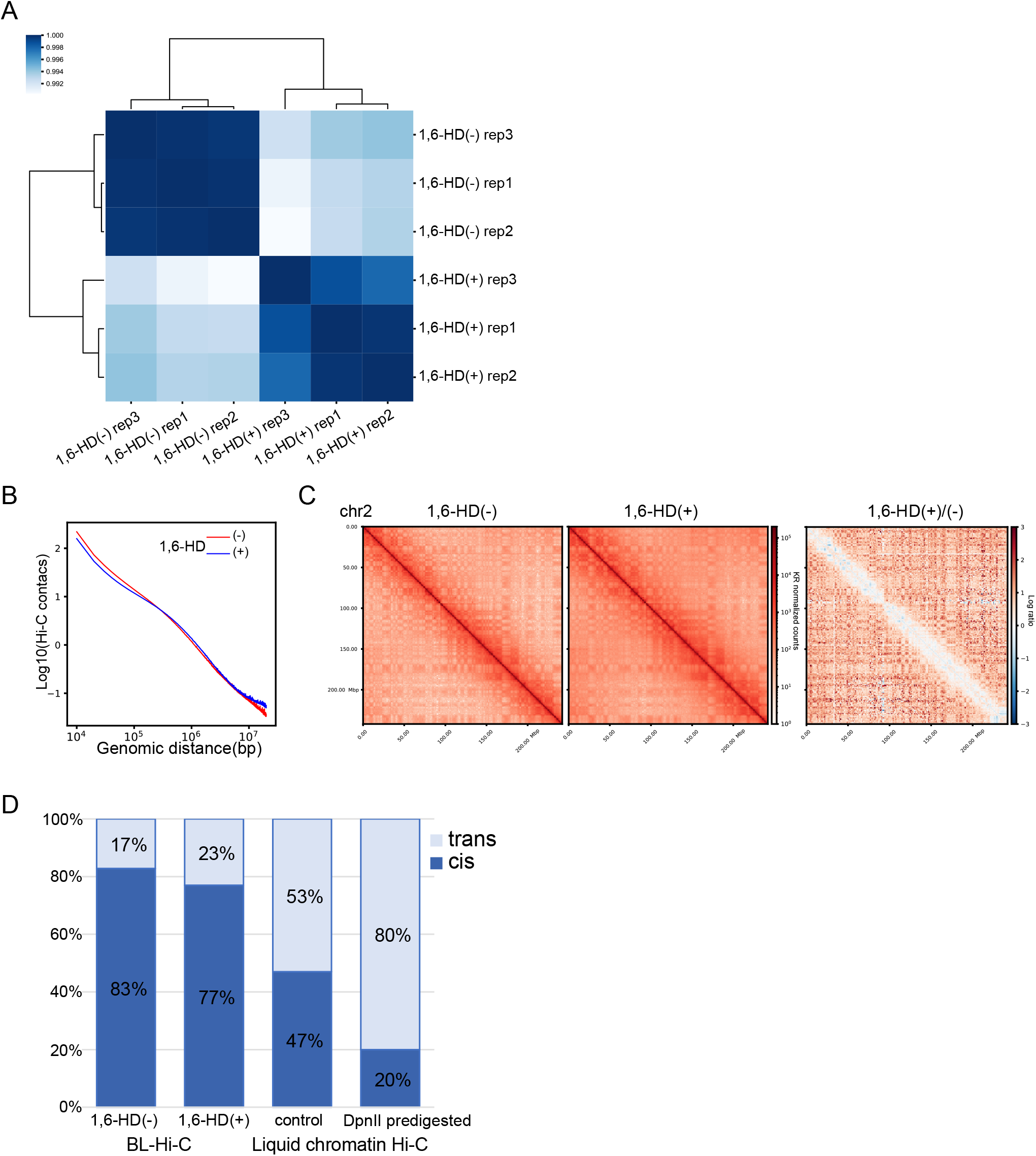
Hi-C interaction changes after 1,6-HD treatment. **A.** Clustering of replicated BL-Hi-C samples. **B.** Genome-wide Hi-C interaction frequency changes before (−) and after (+)1,6-HD treatment. **C.** BL-Hi-C interaction matrices for chr2 at 1Mb resolution. **D.** Data quality of BL-Hi-C and liquid chromatin Hi-C in K562 cells (Belaghzal et al., 2019). Cis: intra-chromosomal interaction, trans: inter-chromosomal interaction. Trans interactions was generally considered to be the noise.

**Figure S4.**
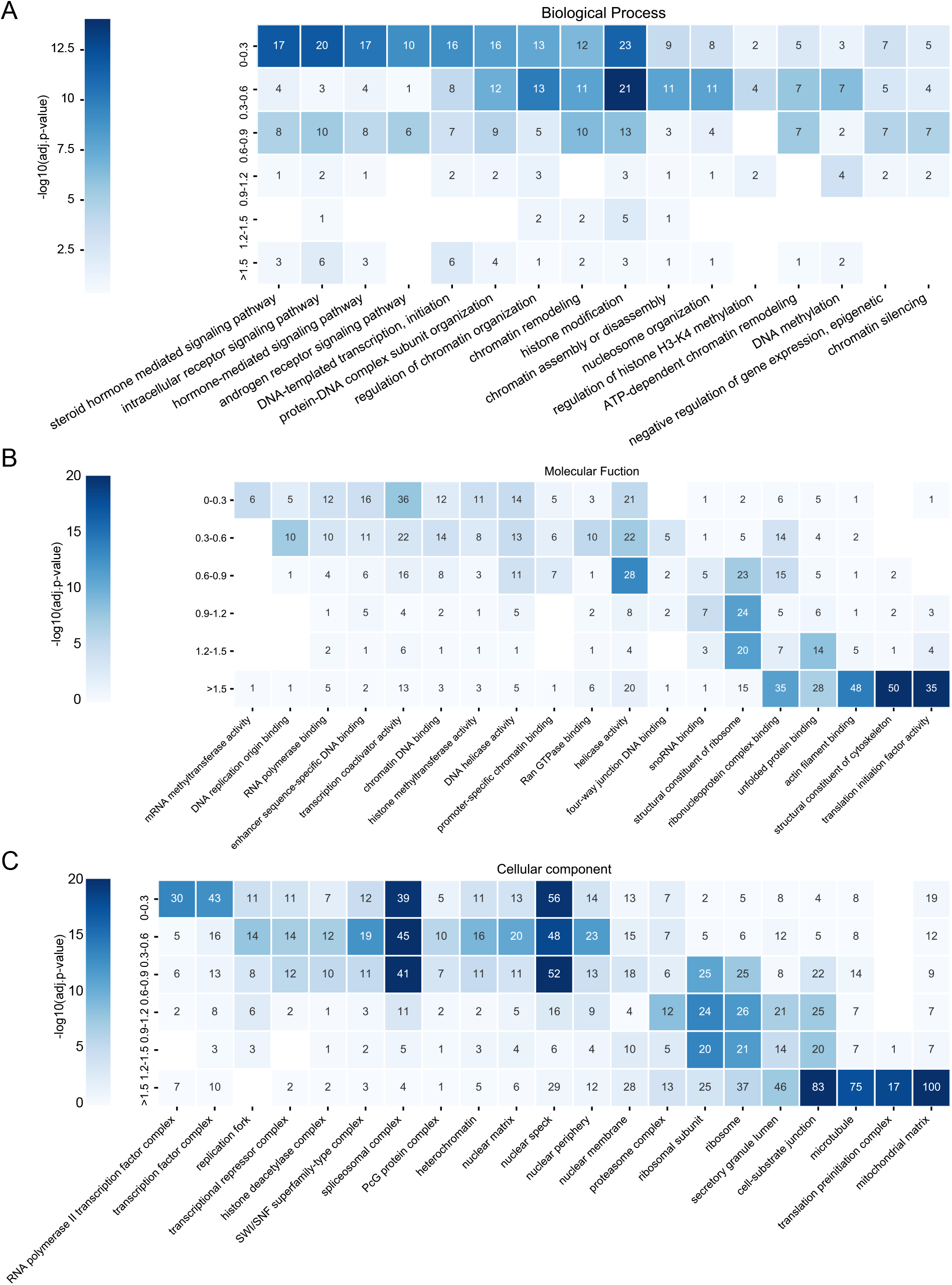
Gene ontology enrichment analysis of proteins. **A.** Gene ontology biological process enrichment analysis of TFs in each group. **B.** and **C.** Gene ontology molecular function (B) and cellular component (C) enrichment analysis of all captured proteins in each group.

**Figure S5.**
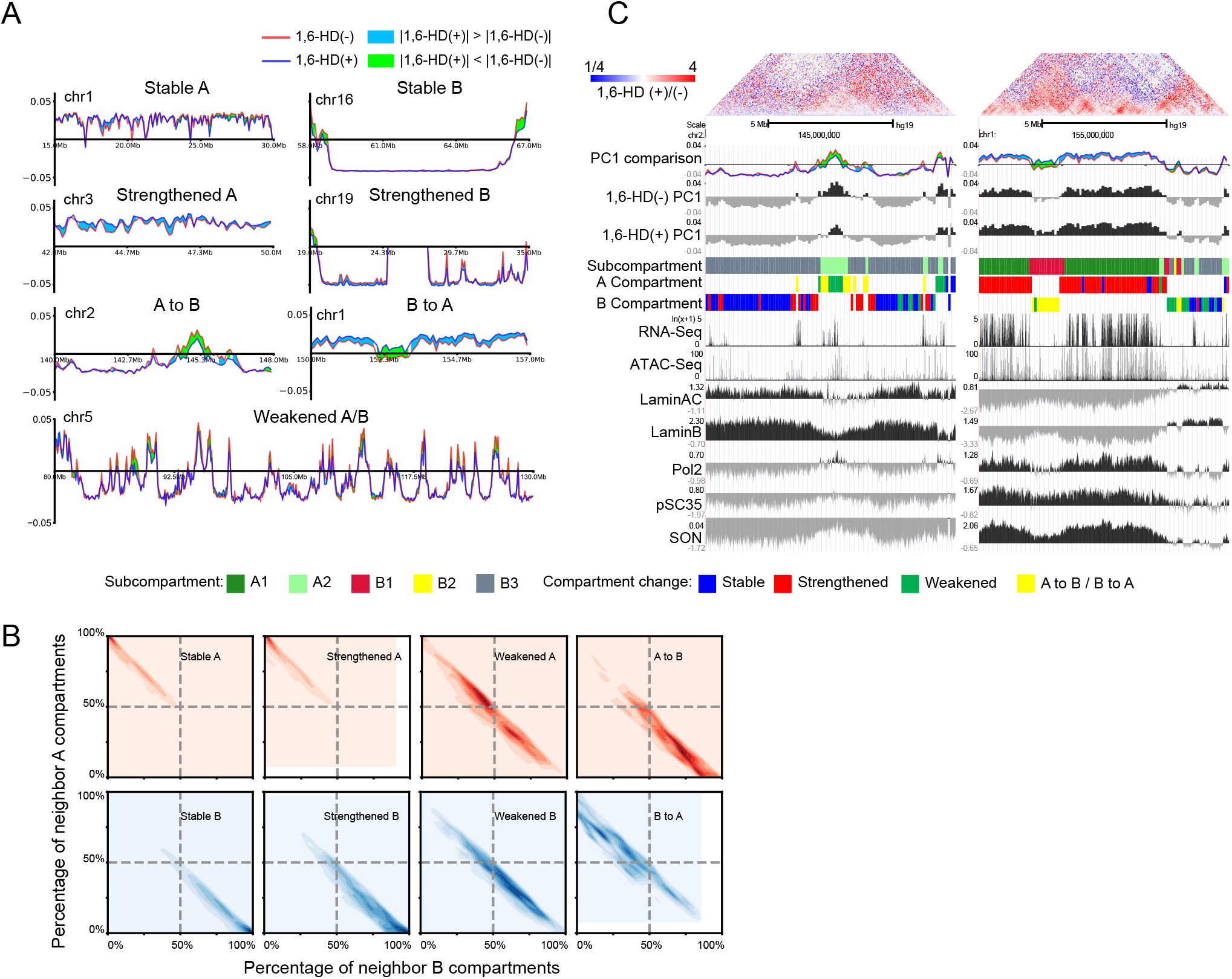
Compartment change-types and neighborhood relationship. **A.** Examples of stable, strengthened, weakened and flipped compartments before (−) and after (+)1,6-HD treatment. The y axis shows PC1 values. **B.** Percentage of neighbor compartments of each compartment change-types. Neighborhood was defined as compartments within 3Mb as usual. **C.** Examples of weakened/flipped A(left) and B(right) compartments and corresponding nuclear speckle/lamina TSA-seq plots.

**Figure S6.**
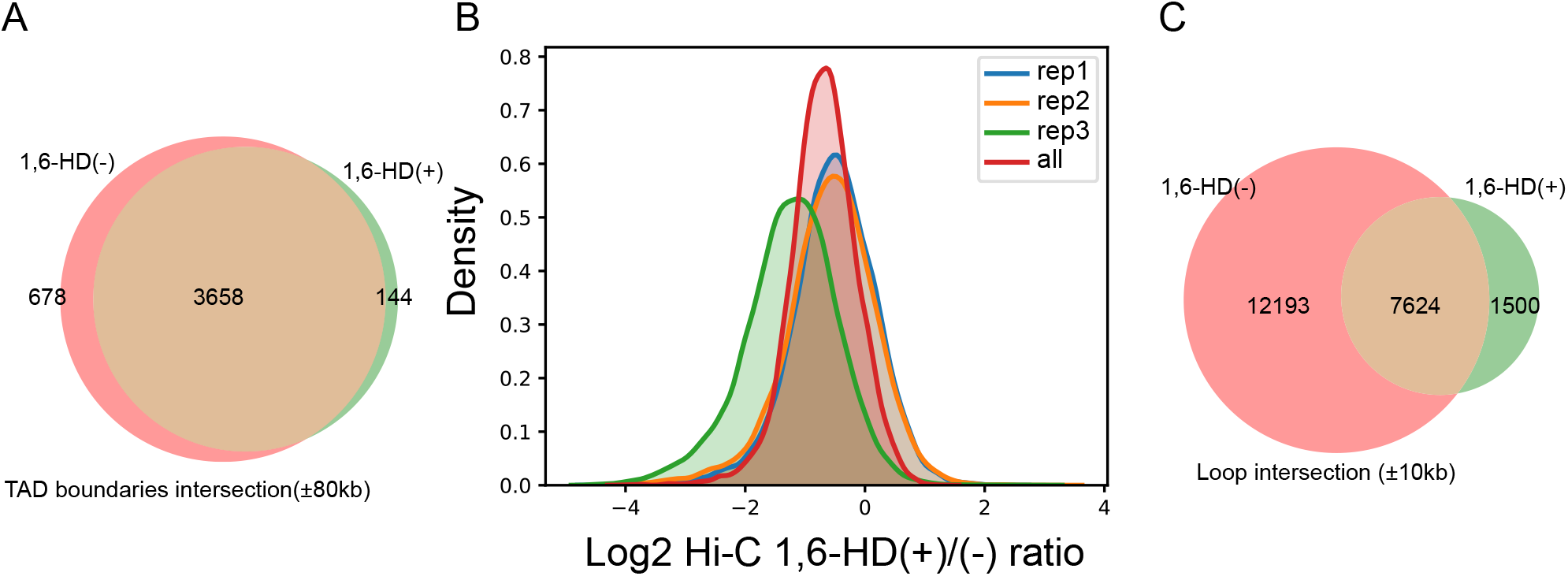
Chromatin TAD and loop changes after 1,6-HD treatment. **A.** Identified TAD boundaries before (−)and after (+)1,6-HD treatment. **B.** DNA loop anchor interaction changes after 1,6-HD treatment. **C.** Identified DNA loops before (−) and after (+)1,6-HD treatment. More than 60% of loops can not be identified again after 1,6-HD treatment.

